# AngioPlate – Biofabrication of perfusable complex tissues in multi-well plates with 4D subtractive manufacturing

**DOI:** 10.1101/2021.08.13.456244

**Authors:** Shravanthi Rajasekar, Dawn S. Y. Lin, Feng Zhang, Alexander Sotra, Alex Boshart, Sergi Clotet-Freixas, Amy Liu, Jeremy A. Hirota, Shinichiro Ogawa, Ana Konvalinka, Boyang Zhang

## Abstract

Organ-on-a-chip systems that recapitulate tissue-level functions have been proposed to improve in vitro–in vivo correlation in drug development. Significant progress has been made to control the cellular microenvironment with mechanical stimulation and fluid flow. However, it has been challenging to introduce complex 3D tissue structures due to the physical constraints of microfluidic channels or membranes in organ-on-a-chip systems. Although this problem could be addressed with the integration of 3D bioprinting, it is not an easy task because the two technologies have fundamentally different fabrication processes. Inspired by 4D bioprinting, we develop a 4D subtractive manufacturing technique where a flexible sacrificial material can be patterned on a 2D surface, change shape when exposed to aqueous hydrogel, and subsequently degrade to produce perfusable networks in a natural hydrogel matrix that can be populated with cells. The technique is applied to fabricate organ-specific vascular networks, vascularized kidney proximal tubules, and terminal lung alveoli in a customized 384-well plate and then further scaled to a 24-well plate format to make a large vascular network, vascularized liver tissues, and for integration with ultrasound imaging. This biofabrication method eliminates the physical constraints in organ-on-a-chip systems to incorporate complex ready-to-perfuse tissue structures in an open-well design.

## Introduction

To address the limitation in existing preclinical models, there is an increasing effort towards building advanced 3D human tissues by incorporating multiple cell types, scaffolds, and dynamic forces. Advances in biofabrication are an integral part of this effort. 3D bioprinting techniques, such as Freeform Reversible Embedding of Suspended Hydrogels (FRESH)^1^ and stereolithography^2^, have been used to build tissues with boundless structural complexity. However, although the tissue production process is generally automated, each 3D printed tissue construct needs to be manually connected for media perfusion or cell seeding before tissue culture can be initiated^3^. This represents a significant engineering manipulation step to transition from tissue printing to perfusion culture. On the contrary, an organ-on-a-chip system based on microfluidic devices or microtiter plates provides built-in perfusion connections^4,5^, but lacks the flexibility of introducing complex 3D tissue structures that are possible with 3D bioprinting^6,7^. Although many key functional features of complex organ systems can be replicated with organ-on-a-chip systems, the physical constraints of plastic microchannels and membranes in these closed microfluidic devices limit the structural design of the tissue microenvironment that can be engineered. To integrate perfusion and complex 3D tissue structures, microfluidic chips could be pre-fabricated, and perfusable networks can be subsequently introduced in the chip with a 3D laser beam microdissection technique^8-10^. While this technique is powerful and produces high-resolution features, it can be slow and difficult to scale. Hence, producing tissues with unrestrained 3D structures, built-in connection for perfusion, and high scalability is not trivial and still much needed.

4D printing, defined as “3D printing + time”, is an emerging concept where the shape of a 3D printed structure can change as a function of time. For instance, encoded with localized swelling behavior, printed composite hydrogels can be programmed to build complex architectures that change shape over time when immersed in water^11^. We exploited this observation and further extended this concept to demonstrate that a flexible sacrificial material patterned on a 2D surface can change shape in 3D when in contact with a natural hydrogel solution. As the gel cross-links, the structurally transformed flexible sacrificial material is locked in place and can subsequently be degraded to produce perfusable networks, which are then populated with human cells to emulate complex organ-specific structures. We termed this method 4D subtractive manufacturing due to the shape changing property of a sacrificial material. As a result, the sacrificial material initially patterned in 2D could easily integrate with a pre-designed microfluidic-based perfusion system and give rise to a 3D structure when triggered at a later stage. Furthermore, taking advantage of the scalability of 2D patterning, we adapted the technique to a format of a 384-well plate (here referred to as AngioPlate™, **Figure 1**) on which we can readily transition from tissue fabrication to perfusion culture for an array of units.

**Figure 1.**
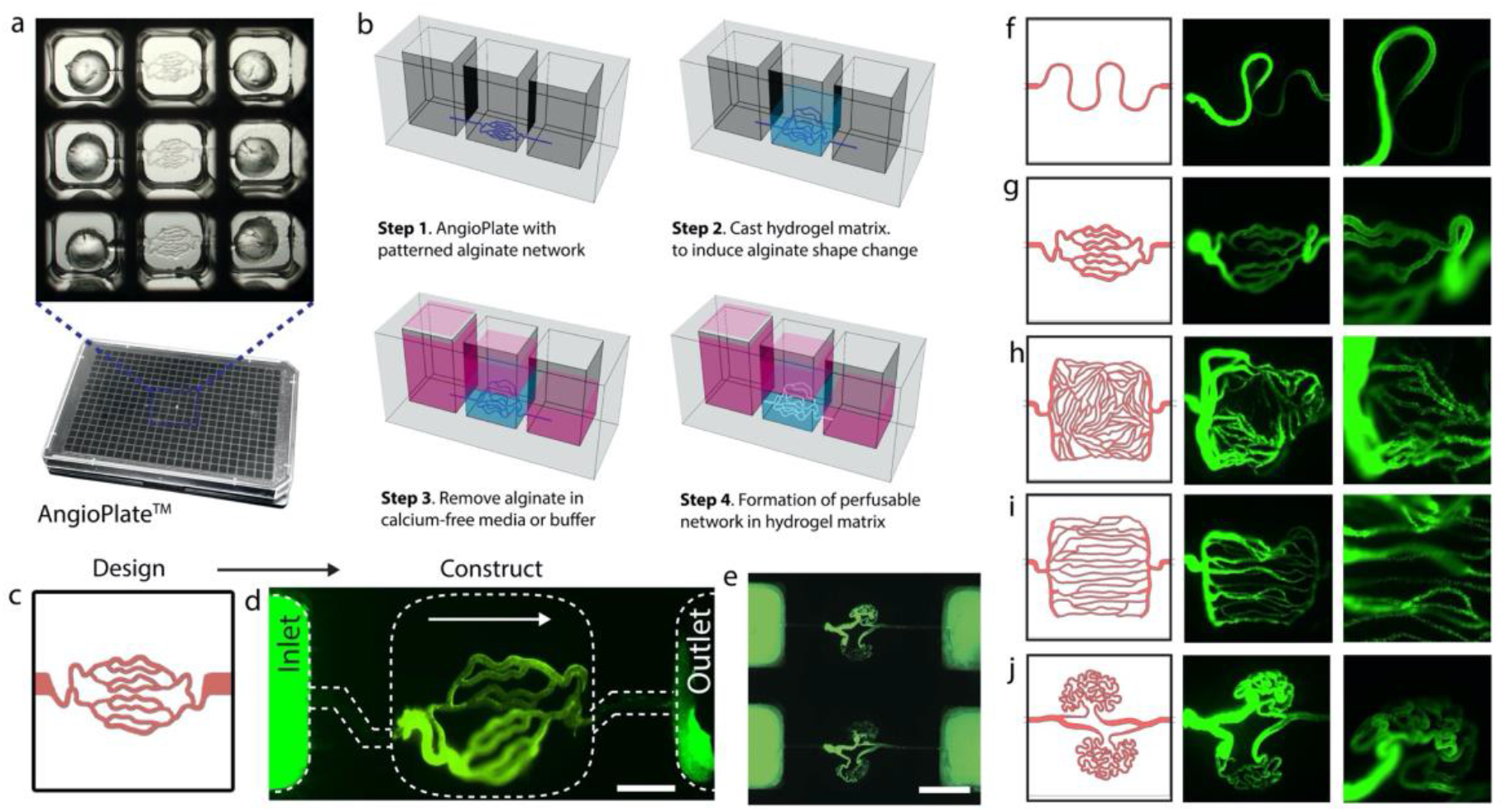
4D subtractive manufacturing and AngioPlate™. **a**, Image of encapsulated alginate fiber networks in a 384-well plate. **b**, Schematic on the use of AngioPlate™, which includes gel casting, folding of the alginate network in the hydrogel, and degradation of the sacrificial alginate template. **c**, Illustration of a generic bifurcating network design. **d**, Perfusion fluorescent particles (1 µm, green) through the assembled 3D network in a hydrogel matrix based on the generic bifurcation network design shown in c. Scale bar, 1 mm. **e**, Perfusion of multiple networks in AngioPlate™. Scale bar, 2 mm. **f-h**, Image of organ-specific networks with structural variation from different initial designs shown on the left and the resulting 3D network perfused with fluorescent particles (1 µm, green) for visualization on the right.

## Results

Our objective is to develop a scalable 3D tissue culture platform (AngioPlate™) that includes built-in fluid connections as in most organ-on-a-chip systems but also a way to obtain tissues with a high level of structural complexity as made possible with 3D bioprinting. To achieve this, we proposed using a flexible sacrificial material that can be patterned in 2D so that it can be integrated into a micro-channel based device for perfusion but can also transform in 3D to produce complex structures when triggered by hydration. The flexible sacrificial material should be inert, patternable, and compatible with various natural hydrogel matrices. Most importantly, it should also degrade in response to a second trigger to carve out a 3D network inside a hydrogel. In search of a material that meets these criteria, we converged on a well-known biomaterial, alginate, which can be rapidly cross-linked and degraded in response to calcium ions. Alginate has been widely used in biological applications as tissue scaffolds^12^ and drug delivery carriers^13^. Using a combination of techniques, including standard photolithography and diffusion-based calcium gelation (**Supplementary Figure 1**), we patterned an array of 128 networks of branched alginate fibers on a polystyrene sheet corresponding to the format of a 384-well plate. The patterned alginate fibers were dried first (**Supplementary Figure 1**) and then assembled against a bottomless 384-well plate (**Figure 1a**). In this format, the plate can be packaged, sterilized with gamma radiation, and then stored.

When using the plate, 25 µL of hydrogel solution (Fibrin or Collagen) were dispensed onto the alginate network to rehydrate the alginate (**Figure 1b, step 1**). During incubation, the dried alginate network quickly swelled, detached from the polystyrene base, and changed shape inside the hydrogel (**Supplementary Video 1**). Then the hydrogel is cross-linked at 37°C (e.g., Collagen and Matrigel) or through enzymatic reaction (e.g., fibrin), locking the folded alginate structure in place (**Figure 1b, step 2**). Finally, phosphate buffered saline (D-PBS) or culture media with low calcium and magnesium concentration (Ca^2+^<200mg/mL and Mg^2+^<100mg/mL) was used to extract the calcium from the alginate and dissolve away the alginate network, resulting in an open perfusable network that is pre-designed to connect with the inlet/outlet wells for perfusion (**Figure 1b, step 3**). Perfusion was initiated with gravity-driven flow by simply tilting the plate at a 15° angle on a programmable rocker (**Figure 1b, step 4**). To ensure the stability of the tissue model over time, the fibrin gel used was highly adhesive and can attach to the polystyrene surface to form a tight seal that prevents leakage. With 1% aprotinin added fresh every other day, fibrin gel degradation was completely prevented. The hydrogel scaffolding was stable, even in the presence of fibroblasts. This feature of fibrin gels is important when compared to collagen, where gel compaction and degradation can be a major issue in long-term culture. However, extra care is needed when working with fibrin gel as the cross-linking speed needs to be fine-tuned by controlling the concentration of thrombin used (**Supplementary Figure 2**).

The extent of alginate folding is determined by the cross-linking speed and viscosity of the hydrogel solution and can result in completely distinct structures from the same initial design, which adds one more dimension of control. However, even under the same gelling conditions, the exact positioning of resulting networks will vary in the 3D space (**Figure 1c-e, Supplementary Figure 3**). In the native tissue, no two biological structures are identical. Hence, this degree of stochasticity is natural. Despite the stochastic behavior, distinct vessel network organizations that resemble various organs or even various parts of an organ can be captured (**Figure 1f-j**). The overall architectural design (i.e., the diameter, density, and location of the branches, etc.) was pre-defined in the initial design (**Supplementary Figure 3**). We have built a 3D network resembling a convoluted tubule (**Figure 1f**), an intricately folded glomerular vessel in a kidney (**Figure 1h)**, densely packed vessels in a liver (**Figure 1i**), and well-aligned vessels as in a muscle (**Figure 1j)**. Although a significant structural transformation from the initial design is not always necessary, the detachment of alginate from the plastic base due to alginate swelling is useful to create a softer microenvironment that is away from direct contact with the hard plastic for seeded cells.

We next populated the network in the vasculature design with endothelial cells which formed a confluent endothelium in 8 days (**Figure 2a**). In some areas, vascular sprouting was also observed (**Supplementary Figure 3**). The resulting vasculature is a 3D structure that contains a hollow lumen and spans the depth of the hydrogel as shown from the confocal x-z plane view (**Figure 2a and 2e**). The vascular network in the presence of a confluent endothelium was significantly less permeable and displayed barrier function to large proteins. When exposed to Triton™ X-100 and TNF-α, the vascular permeability increased significantly and showed a dose dependent response to TNF-α. When exposed to lipopolysaccharides (LPS) from E. coli to simulate infection, the networks showed no significant changes in permeability and barrier function (**Figure 2b-c**). The endothelial cells also deposited a basement membrane that contained laminin which was localized on the basal side of the endothelium (**Figure 2d**). Furthermore, the endothelial cells produced and distributed von Willebrand factor (vWF) both intracellularly and extracellularly, which is a key protein for regulating blood coagulation (**Figure 2f**). Lastly, transmission electron microscopy showed that the endothelial cells form intercellular tight junctions consistent with the vessel barrier function shown (**Figure 2g**).

**Figure 2.**
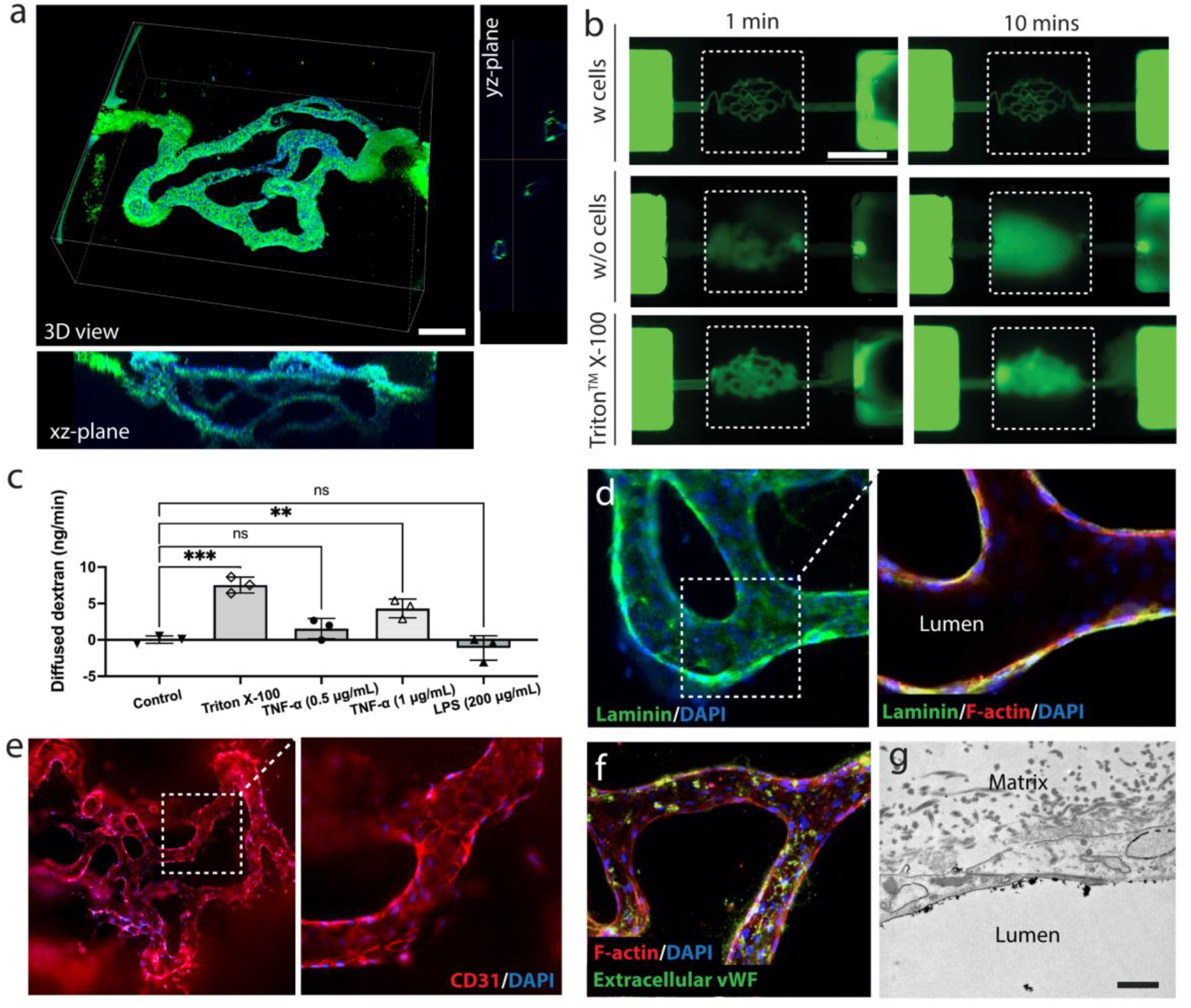
AngioPlate™ vasculature. **a**, 3D volumetric rendering of confocal z-stack images of a vascular network populated with human umbilical cord vein endothelial cells. View from xz-plane is maximum projection. Cells were stained for CD31 (green) and DAPI (blue). Scale bar, 500 µm. **b**, Fluorescent images of vascular networks perfused with 70kDa fluorescein-dextran (green) with or without the presence of endothelial cells and/or Triton™ X-100 treatments. Scale bar, 2 mm. **c**, Quantification of dextran (70kDa) diffusion rates of the vascular networks in response to no treatment, Triton™ X-100 (positive control), and drug treatments, n=3. Statistical significance was determined using one-way ANOVA. ns indicates not significantly different. *P ≤0.05 **P≤0.01 ***P ≤0.001. **d-f**, Fluorescent image of a vascular network stained for Laminin (green), F-actin (red), CD31(red), extracellular vWF (green), and DAPI (blue). **g**, Transmission electron microscope image of endothelium on a vessel wall showing the tight junction between two adjacent endothelial cells. Scale bar,1 µm.

Expanding the 4D subtractive manufacturing method, we incorporated multiple individually perfusable networks within the same matrix to reproduce the spatially intertwined vascular-tubular networks that emulate the vascularized proximal tubule complexes in a kidney (**Supplementary Figure 4a-d**), as well as terminal lung alveoli composed of both alveolar duct and sac components (**Supplementary Figure 4e-c**). For the kidney model, we have three networks perfused with three inlet and three outlet wells (**Figure 3a**). The networks were designed to be in proximity to each other as much as possible to facilitate mass transport (**Figure 3b-c**). The proximal tubule cells formed a confluent epithelium that tends to be thicker than the endothelium barrier (**Figure 3d-e**). The epithelium contains primary cilia labeled by α-Tubulin (**Figure 3f**), glucose transporters (**Figure 3g)**, and Na^+^/K^+^-ATPase (**Figure 3h)**, which are key transport proteins. We also found that the epithelial cells can deposit laminin which forms the basement membrane on the basal side of the epithelium (**Figure 3i)**. The tissues can also be extracted and sectioned for immunohistochemistry (**Figure 3j**). Both the tubular and the vascular compartments maintained high barrier function that can confine large proteins (**Figure 3k-l**). By day 14 post confluency, a glucose gradient between the tubule and the surrounding vasculature can be established 24 hours after glucose injection into all networks indicating the presence of glucose reabsorption function in the tissue (**Figure 3m**).

**Figure 3.**
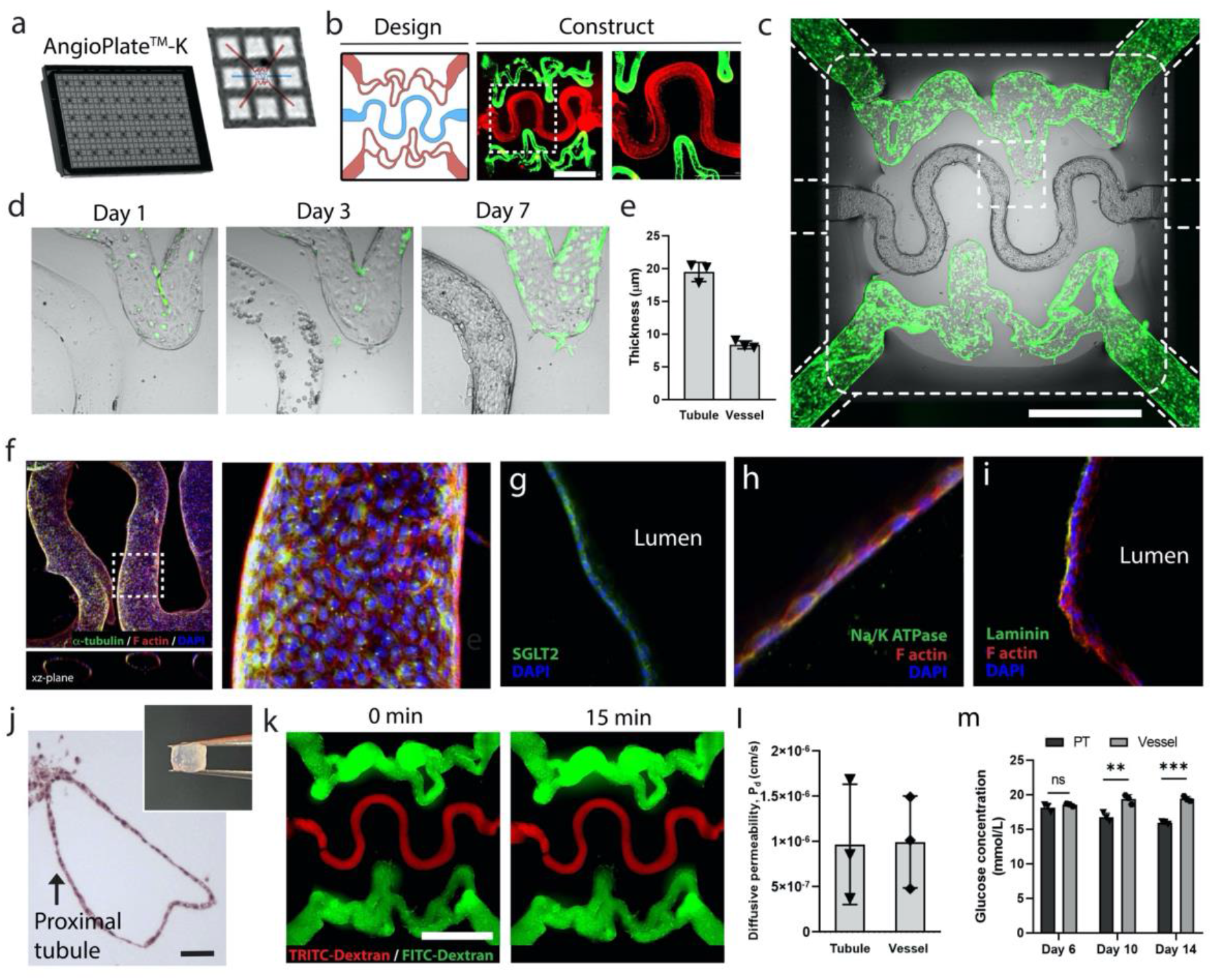
AngioPlate™ vascularized proximal tubule complex. **a**, Configuration of AngioPlate that allocates three inlet wells and three outlet wells for each kidney model. **b**, Design of the vascular proximal tubule complex and the resulting acellular network perfused with FITC particles (1 µm, green) and TRITC particles (1 µm, red) for visualization. Scale bar, 1 mm. **c**, AngioPlate capillary-proximal tubule complex seeded with GFP-HUVECs (green) and kidney proximal tubule cells on day 7. Scale bar, 1 mm. **d**, Zoomed-in time-lapse images of capillary-proximal tubule complex throughout the cell seeding process. **e**, Quantification of the thickness of epithelium and endothelium on day 7 (n=3). **f**, Confocal image of a proximal tubule stained for F-actin (red), α-tubulin (green), and DAPI (blue). **g-i**, Fluorescent images showing the cross-section of the proximal tubule epithelium stained for F-actin (red), SGLT2 (green), Na/K ATPase (green), Laminin (green), and DAPI (blue). **j**, Histology tissue section of a proximal tubule stained with Masson’s trichrome. Insert showing a fixed whole tissue removed from the AngioPlate™ ready for histological sectioning. **k**, Capillary-proximal tubule complex perfused with TRITC-Dextran (red, 70kDa) in the tubule and FITC-Dextran (green, 70kDa) in the vascular network. Perfused networks imaged at 15 minute intervals. Scale bar, 1 mm. **l**, Quantification of diffusive permeability to 70kDa dextran, n=3. **m**, Glucose levels in media perfusate from vascular and tubular networks on days 6,10 and 14 (n=3). Proximal tubule cells were seeded on day 4. Statistical significance was determined using one-way ANOVA with the Holm-Sidak method. *P≤0.05 **P ≤0.01 ***P ≤0.001

To model inflammatory disease condition in the kidney model, the tissues were stimulated with TNF-α. TNF-α was perfused through the vasculature for 12 hours (**Figure 4a**). The expression of ICAM-1, a surface glycoprotein important for leukocyte recruitment, on the endothelium was significantly increased due to TNF-α treatment (**Figure 4b-c**). The perfusates from both the vascular and tubular compartments were collected and analyzed for a range of cytokines secreted by the cells. We saw a significant increase in IL-8, MCP-1, GM-CSF production due to TNF-α stimulation (**Figure 4d**). MCP-1 and GM-CSF are responsible for recruiting circulating monocytes which is a typical process that takes place during tissue inflammation. IL-6 showed an increasing trend. No significant changes were observed in the other 11 cytokines analyzed (**Supplementary Figure 5**). Although a strong effect was observed in the vascular compartment, the effect on the kidney tubule is relatively mild as most TNF-α delivered through the vasculature was confined within the vasculature within the 12-hour treatment period. This indicates the source of inflammation (vascular circulation or tissue parenchymal) will have an effect on tissue response, at least in the short term. We did, however, suspect there is occasional leakage in the vascular or the tubular compartment due to injury which can lead to a different cytokine profile in the tubular compartment as shown by the outlier data points in TNF-α, IL-8, GM-CSF secretion (**Figure 4d**).

**Figure 4.**
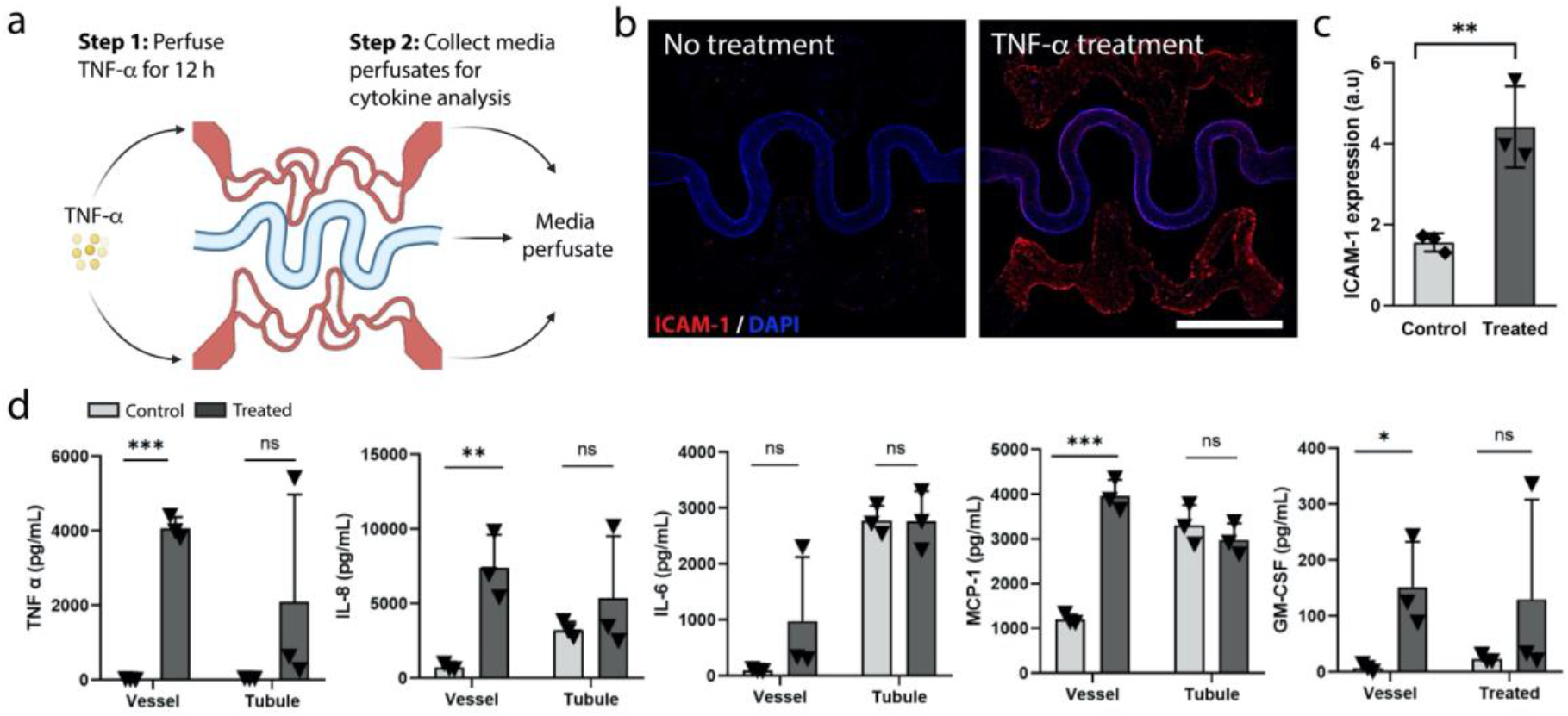
Cytokine secretion analysis of AngioPlate™ vascularized proximal tubule complex under inflammation. **a**, Experimental setup for TNF-α stimulation and sample collection for cytokine analysis. **b**, Tissues stained for ICAM-1 (red) and DAPI (blue) after TNF-α stimulation. Scale bar, 1 mm. **c**, ICAM-1 expression levels quantified using ImageJ, n=3. **d**, Quantification of cytokine levels in media perfusates, n=3. Statistical significance was determined using one-way ANOVA and one-way ANOVA on ranks with the Holm-Sidak method. ns indicates not significantly different. *P≤0.05 **P≤0.01 ***P ≤0.001

To develop a vascularized terminal lung alveoli model, three inlet and two outlet wells were used (**Figure 5a**). We developed a structure resembling the terminal lung alveoli with an alveolar duct and five alveolar sacs with no outlets (**Figure 5b**). Because the fibrin gel is porous, cells can still be perfused into the network as liquid escapes the network via interstitial flow through the porous gel. In 4 days, the alveolar epithelial cells can coat the entire network (**Figure 5c-e**). From day 4 to day 7, we found the epithelium thickness significantly increased, a sign of maturation (**Figure 5e**). However, because the cell source is a cancerous alveolar cell line (A549), the cells have a tendency to overpopulate and physically obstruct the entire chamber around one week after confluency. Therefore, there was a narrow time window for experimental work. A different cell line or healthy primary cells might be needed to establish a more stable model. Nonetheless, on day 7, the alveolar epithelium barrier becomes polarized as indicated by the stronger F-actin staining developing on the apical side of the epithelium (**Figure 5f**). The lung alveolar epithelium is in close proximity to the vasculature where in some locations the two structures physically touched each other but maintained distinctly separated (**Figure 5g**). Histology tissue sections showed the presence of intercellular junctions as visualized by E-Cadherin staining as well as CD31 positive vascular networks in close proximities to the alveoli sac (**Figure 5h**).

**Figure 5.**
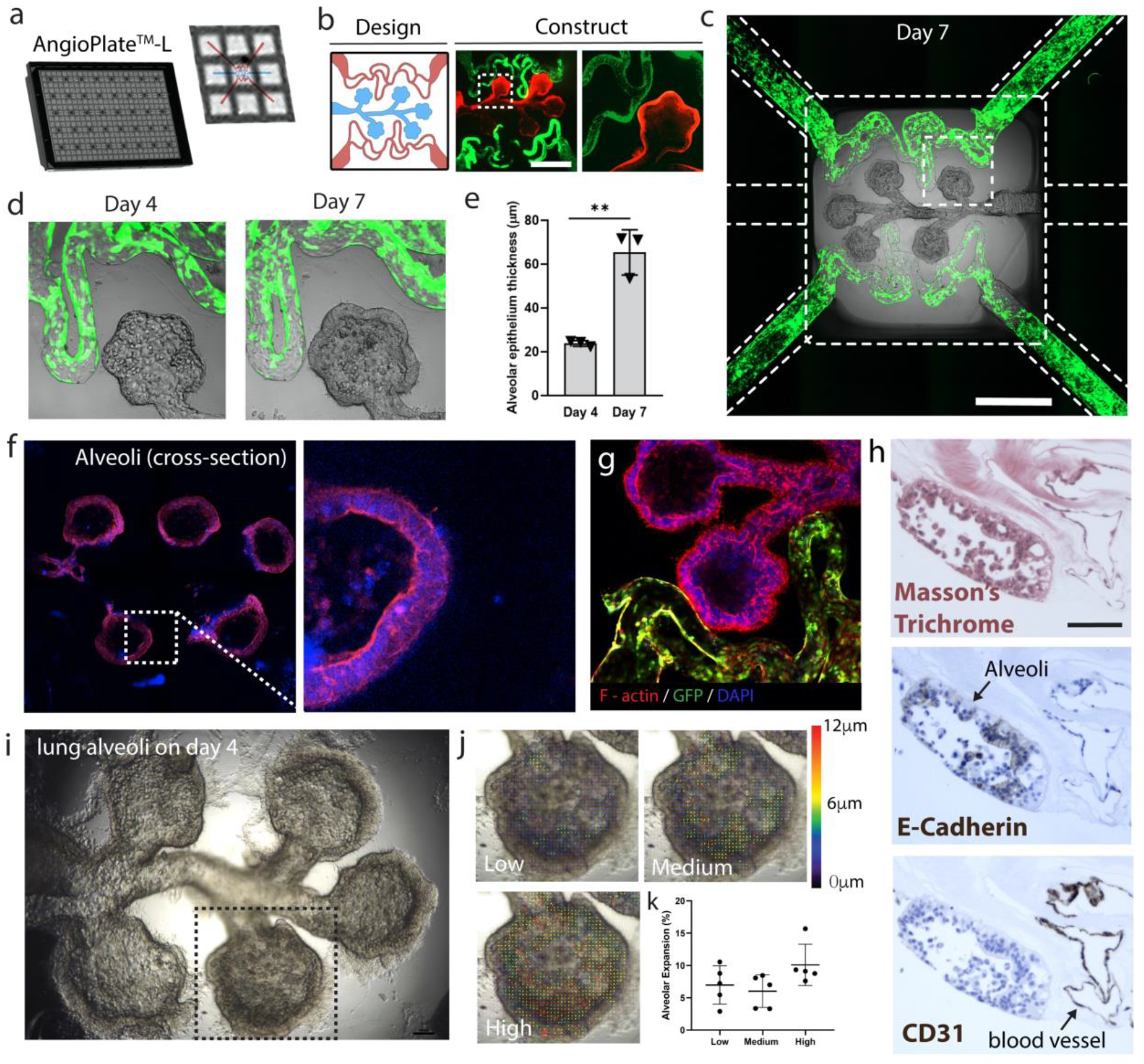
AngioPlate™ vascularized terminal lung alveoli. **a**, Configuration of AngioPlate that allocates three inlet wells and two outlet wells for each terminal lung alveoli model. **b**, Design of the vascularized alveoli terminal with alveolar duct and sac and the resulting network perfused with FITC microparticles (1 µm, green) and TRITC microparticles (1 µm, red) for visualization. Scale bar, 1 mm. **c**, Vascularized alveoli terminal seeded with GFP-HUVECs (green) and alveolar epithelial cells (A549) on day 7. Scale bar, 1mm. **d-e**, Image and quantification of alveolar epithelium thickening over time. **f-g**, Fluorescent images of the cross-section of the vascularized alveolar sac stained for F-actin (red) and DAPI (blue). **h**, Histology cross-section of an vascularized terminal alveoli stained with Masson’s trichrome, E-Cadherin and CD-31. scale bar, 100 µm. **i**, A frame of a video that shows the physical expansion of the alveoli terminal actuated by a ventilator. **j-k**, Vector map and quantification that show the physical expansion of an alveolar sac at different levels of ventilation pressure. Statistical significance was determined using one-way ANOVA with the Holm-Sidak method. *P≤0.05 **P≤0.01 ***P ≤0.001

To demonstrate that the lung structure can be mechanically induced to simulate breathing, we developed a customized lid that can distribute air to the inlet well of the lung structure (**Supplementary Figure 6**). The lid was connected to a ventilator that can repeatedly pump air in and out of the wells. In response to changing air pressure, we observed the lung alveolar duct and sac could physically expand (**Figure 5i**). The extent of the expansion increases with increasing pressure inputs (**Figure 5j-k, Supplementary Video 2**). This demonstration indicates that complex 3D branched structures without any outlets can also be created using our 4D subtractive manufacturing technique and the built-in perfusion connection is robust even to withstand mechanical stimulation. However, removing liquid from this 3D structure to establish an air-liquid interface was challenging and will require further optimization. But recent studies showed that lung epithelium could also maturate under immersion without the air-liquid interface^14^, which could be a better approach for this model.

To demonstrate that the 4D subtractive manufacturing method can be scaled to make larger tissues, which is important for therapeutic applications, clinical imaging or applications that require surgical tissue manipulations, we applied the technique to a 24-well plate (AngioPlate™-24) with a larger well size (15.6mm in diameter, **Figure 6a**). A large branched vascular network was produced using the same procedure (**Figure 6b-c**). Endothelial cells cultured within the network formed a tight vascular barrier and showed signs of vascular sprouting starting on day 14 (**Figure 6d**). Solid parenchymal tissues, such as hepatic spheroids, can also be incorporated within the matrix around the vasculature to make a vascularized liver tissue (**Figure 6e**). Albumin was produced by the hepatic spheroids for at least 20 days and can be collected through the built-in vasculature from the inlet and outlet wells (**Figure 6f**). The tissue can then be extracted from the well for implantation applications while preserving the tissue structures (**Figure 6g**). The larger well also allows easy integration with clinical bioimaging methods such as photoacoustic imaging. An ultrasound gel can be placed on top of the tissue followed by an ultrasound probe. A contrast agent IR-783 dye used for ultrasound imaging was perfused through the vasculature and the perfused vessel network was visualized (**Figure 6 h-i**). Although the imaging resolution is lower than standard fluorescent imaging, it allowed much deeper sample penetration from 5 cm away. The entire imaging path from the probe to the sample is filled with a hydrogel that emulates the density of human tissues to avoid any ultrasound signal disruption. Photoacoustic actuation could potentially be applied in the future to model ultrasound-triggered drug delivery *in vitro*.

**Figure 6.**
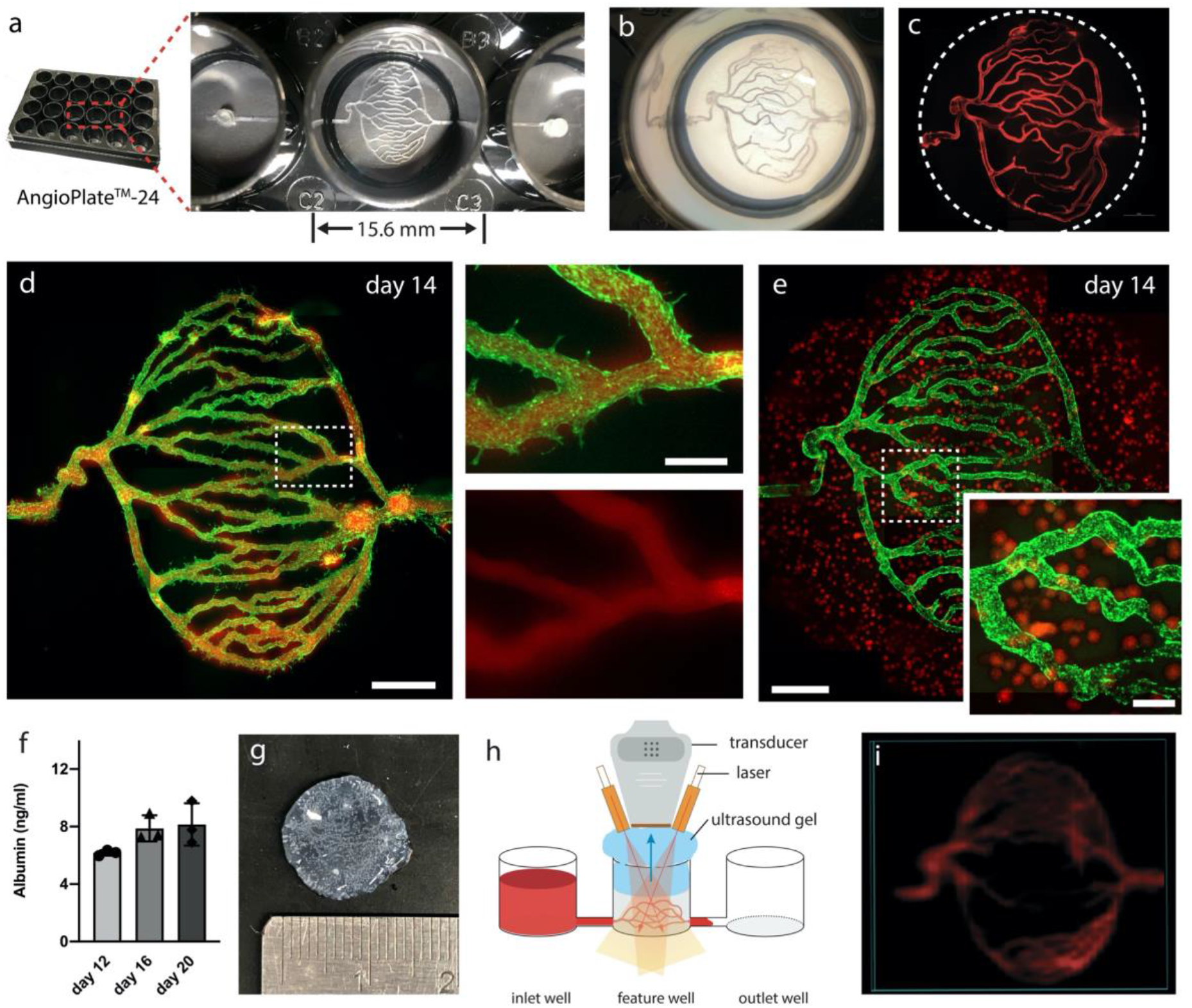
Large AngioPlate vasculature for tissue vascularization and integration with photoacoustic imaging. **a**, Configuration of AngioPlate for producing large vascular networks based on a 24-well plate format. **b**, Image of a vascular network inside a well of an AngioPlate™-24. **c**, Fluorescent image of the network perfused with TRITC particles (1 µm, red) for visualization. **d**, Fluorescent image of a vascular network coated with GFP-HUVECs on day 14. Insets show one area of the network perfused with 70kDa TRITC-dextran (red). Scale bar, 2 mm. Scale bar for inserts: 500 µm. **e**, Fluorescent image of a vascularized liver tissue co-cultured with GFP-HUVECs and hepatic spheroids on day 14. Scale bar, 2 mm. Scale bar for inserts, 500 µm. **f**, Quantification of albumin production over time, n=3. **g**, Image of an extracted vascularized liver tissue from AngioPlate™-24. **h**, Illustration of the photoacoustic imaging setup. **i**, Photoacoustic image of the vascular network perfused with IR-783 dye.

## Discussion

The 4D subtractive manufacturing technique not only takes advantage of the scalability of 2D patterning to integrate with widely-adopted multi-well plates for ease of use but also applies a shape-changing sacrificial material to transform a 2D pattern into a 3D perfusable structure for improved model complexity. Although the manufacturing of the device involves multiple steps, the platform streamlines the process from tissue production to perfusion culture without compromising experimental throughput and model design. Furthermore, the time to build the tissue structure with the current method is completely independent of the complexity of the structures and is usually completed within 15-20 minutes which is the typical duration for matrix gelation. Therefore cells or tissues embedded within the gel matrix will experience minimal stress during the biofabrication process.

Another key feature of the AngioPlate™ is the open-well design. Different from the conventional closed microfluidic-based systems, AngioPlate™ allows easy tissue extraction for downstream analysis. We demonstrated that not only is the platform compatible with histopathological assessment, which is the gold standard in clinical diagnosis of disease and drug injury, but when scaled up, is also compatible with clinical imaging techniques, such as photoacoustic imaging. Molecular analyses including RNA sequencing and proteomics are also possible experimental readouts with this system. These features could allow direct comparison of the *in vitro* data with human clinical data in future studies for model validation.

The open-well design also allows pre-fabricated tissue to be easily added to each well. For instance, a monolayer of organ-specific epithelial cells could be cultured on the top surface of the gel with a supporting perfusable vascular network underneath. Alternatively, tissue spheroid, organoids or tissue explants could be vascularized by placing on top of the gel and the supporting vascular bed. The platform could also potentially be integrated with extrusion-based 3D printing in the future to expand model complexity further. By having an open-well design and by removing the geometric constrain of synthetic membranes or microfluidic channels, the platform becomes highly versatile. The tissue models, built entirely inside a natural hydrogel with no restrictive boundaries, could be seeded with stem cell-derived organoids and serve as the initial structural template to study tissue development and morphogenesis^15^. Lastly, when using the AngioPlate™, users need only pipetting techniques to handle all reagents and the protocol. It does not require users to assemble tubes or pumps, making the platform compatible with the robotic fluid handling system for automation.

## Limitations

Although the AngioPlate™ provides a new approach of constructing complex tissues and establishing tissue perfusion, there are still limitations. From the aspect of device fabrication, our current process uses PDMS glue to bond a bottomless well plate with a polystyrene base. PDMS is prone to drug adsorption^16^ and may need to be replaced depending on the application. In addition, the spread of the PDMS glue into the middle well is not always easy to control, which could damage the patterned features and lead to device failure. This problem is mitigated in larger wells such as in the 24-well plate design. Ultrasound or laser welding methods could also be used to circumvent this issue in device bonding for industrial-scale manufacturing.

From a design aspect, different from 3D printing, our current method does not provide the flexibility of changing the design on demand. However, the master molds we used for alginate patterning could be 3D printed without the time-consuming photolithography step, which would combine the scalability of the 2D patterning with the versatility of 3D printing to allow rapid design iterations. Alternatively, the patterned 2D alginate features could be directly 3D printed with an extrusion-based 3D printer without using any PDMS molds, which could significantly shorten the fabricate process while providing even more design flexibilities.

To avoid the use of any calcium chelating agents that could potentially damage cells, we used buffer or culture media with low calcium ions concentrations for the degradation of the alginate sacrificial material. This degradation is a slower process that can last two days. But considering the overall length of a typical culture and tissue maturation process, this is a minor delay from an end-user perspective. During tissue culture, it’s important to note that media perfusion is entirely driven by gravity which varies over time due to the depletion of the pressure heads. Moreover, the flow pattern is bi-directional. If a constant unidirectional flow is required, customized lids with built-in microfluidic pumps could be used to recirculate the media from outlet wells back to inlet wells to maintain fluid pressure^17^.

## Conclusions

The trade-off between model complexity and experimental throughputs has been a long-standing issue^18^. More studies are needed to understand the level of tissue complexities that are necessary in model systems which is likely specific to the biological question of interests. But when complex tissue structures and the use of natural extracellular matrix are needed, the 4D subtractive manufacturing technique could offer a new way of approaching tissue production and culture to close the existing disparity between model complexity and throughputs. Moreover, the emergence of stem cell-derived organoids will require platform technology that can accommodate organoid growth with an amenable matrix and offer perfusion capability without compromising experimental throughputs^7^. In future work, the integration of organoids and the AngioPlate™ platform could be explored^19^.

## Materials and Methods

### AngioPlate™ fabrication

Standard photolithography techniques were used to create each SU8 master mold. To create a negative mold, a PDMS (Ellsworth Adhesive, Cat# 4019862) mixture at a ratio of 1:30 was poured onto the SU8 master mold and cured at 47 °C overnight. The cured PDMS mold was soaked in 5% w/v pluronic acid (Sigma Aldrich, Cat# P2443) for 30 minutes, washed with distilled water, dried, and then capped onto a plasma-treated polystyrene sheet (11.5 × 7.5 cm, Jerry’s Artarama, Cat# V16013). Assembled PDMS mold and polystyrene sheet were then transferred to a one-well plate. To fill the entire pattern with 3% w/v alginate solution (Sigma Aldrich, Cat# A2033), 75mL of the solution was poured into the one-well plate containing assembled PDMS mold and sheet. After the alginate solution filled up the patterned microchannels, the residual solution was aspirated out. To cross-link alginate, 75mL of 5.5% w/v calcium chloride solution (CaCl_2_) (Sigma Aldrich, Cat#223506) was added and left to cross-link overnight. After cross-linking, the residual CaCl2 solution was removed and rinsed with distilled water. Cross-linked alginate within the PDMS channels was then air-dried for 48 hours. Melted poly (ethylene glycol) dimethyl ether (PEGDM) (Sigma Aldrich, Cat# 445908) was then injected into the channels and allowed to fill at 70 °C for 1 hour. Once PEGDM re-solidifies, PDMS mold was peeled off and the alginate fibers encapsulated in PEGDM were transferred onto the polystyrene sheet. The polystyrene sheet containing the patterns was then bonded onto a bottomless 384-well plate using a high viscosity PDMS glue (Ellsworth Adhesive, Cat# 2137054) at a ratio of 1:10. The assembled device was allowed to cure overnight at room temperature. Assembled devices were sterilized for 2.5 hours using 70% w/v ethanol and PEGDM was washed off during the sterilization process. The plates were air-dried inside a biosafety cabinet (BSC) for 12 hours and stored at 4 °C until use.

### Cell culture

Green Fluorescent Protein-tagged Human Umbilical Vein Endothelial Cells (GFP-HUVECs, Angio-Proteomie, Cat# CAP-0001GFP) were grown in Endothelial Cell Growth Media (ECGM2, (PromoCell, Cat# C-22011) in cell culture flasks coated with 0.2% w/v gelatin (Sigma Aldrich, Cat# G9391). Renal Proximal Tubule Epithelial Cells (RPTEC-TERT1, Everycyte, Cat# CHT-003-0002) and Lung Alveolar Epithelial Cells (A549, Cedarlane Labs, Cat# PTA-6231) were cultured as per manufacturer’s instructions. RPTEC-TERT1 cells were grown using the ready-to-use ProxUp media from Evercyte (Cat# MHT-003). A549 cells were cultured in F-12K media (Cedarlane Labs, Cat# 302004) supplemented with 10% fetal bovine serum (FBS, Wisent Bioproducts, Cat# 098-150), 1% penicillin-streptomycin solution (100X, Wisent Bioproducts, Cat# 450-201-EL) and 1% HEPES (1M, Wisent Bioproducts, Cat# 330-050-EL). Cells were grown in T75-cell culture flasks in an incubator maintained at 37 °C and 5% CO_2_ until they were 80% confluency before being passaged for further experiments. Cells were strained through a 40 µm cell strainer to get a single cell suspension prior to cell seeding. All cells used in this work were between passages 2-6.

Human C3A hepatocellular carcinoma cells (HepG2s, ATCC, Cat# CRL-10741) and primary human lung fibroblasts were used (Cedarlane Labs, Cat# PCS-201-013) to generate hepatic spheroids. HepG2s were cultured in Eagle’s minimal essential medium (EMEM) containing 10% FBS, 1% penicillin-streptomycin and 1% HEPES solution. Primary human lung fibroblasts were cultured in Dulbecco′s Modified Eagle′s Medium (DMEM) containing 10% FBS, 1% penicillin-streptomycin and 1% HEPES solution. HepG2s were labeled with CellTacker Red CMTPX dye (Thermo Fisher Scientific, Cat# C34552) to locate the spheroids following the supplier’s instruction. Aggrewell800™ plates were used to generate hepatic spheroids. 1 mL of cell suspension containing 3 million cells (about 800 HepG2s and 200 lung fibroblasts per spheroid) were added per well. To prevent cell adhesion on the plate, an anti-adherence rinsing solution (STEMCELL technologies, Cat# 07010) was used to treat the Aggrewell800™ plate. The spheroids were cultured in the Aggrewell800™ plate for eight days and then collected for seeding.

### Hydrogel casting and device operation

Hydrogel used for AngioPlate was 10 mg/mL fibrinogen (Sigma Aldrich, Cat# F3879). Before hydrogel casting, the hydrogel was aliquoted into 125 µL aliquots and each aliquot was mixed with 25 μL of thrombin (7 U/mL, Sigma Aldrich, Cat# T6884). Immediately after mixing, 25 μL of gel solution was then cast into each well. The plate was left in a BSC for 30 minutes to allow for gelation. Alternatively, neutralized Rat Tail Collagen (2-2.5mg/mL, Corning, Cat# CACB354249) with an extra 20% v/v D-PBS (Thermo Scientific, Cat# 14190144) can be used as another source of natural extracellular matrix. To degrade the alginate polymers, D-PBS was added to all wells and the platform was placed on a rocker at 4 °C for 48 hours with daily buffer change. Alternatively, DMEM containing 10% FBS, 1% penicillin-streptomycin and 1% HEPES solution was added to all wells and the platform was placed in an incubator at incubated at 37°C and 5% CO_2_ for 48 hours with daily media change. The device was then primed again with cell culture media and incubated at 37°C and 5% CO_2_ for 24 hours prior to cell seeding. After removing all the media, 30 μL of fresh media was added into the middle well and 125 μL of the cell suspension (0.6 million cells/mL) was added to the inlet and outlet wells. The plate was placed flat in the incubator for 2 hours to allow cell attachment. After cell attachment, culture media was removed, and 50 μL fresh media was added to the middle well, and 90 μL to the inlet and outlet wells. To initiate perfusion, the entire platform was placed on a programmable rocker that tilts at a 15° angle and alternates its tilting direction every 15 minutes. 1% v/v aprotinin (2 mg/mL, Sigma Aldrich, Cat# 616370-M) was added to all media to prevent fibrin gel degradation. Media was changed every other day. For the vascularized kidney proximal tubule model, endothelial cells were seeded first and proximal tubule cells were seeded on day 3. For the vascularized alveolar terminal model, endothelial cells were first and the lung alveolar cells were seeded on day 3. For the vascularized liver tissue in AngioPlate™24, each tissue contains about 2400 hepatic spheroids. The hepatic spheroids were pre-mixed with fibrin gel and then cast into the plate. Alginate was degraded with DMEM media inside the incubator and endothelial cells were seeded after alginate degradation. DMEM and ECGM2 media at a ratio of 1:1 (v/v) containing 1% v/v Aprotinin were used as the co-culture media and the media were changed daily. To test alginate degradation, the networks were perfused with fluorescent FITC and/or TRITC latex beads (1.0 µm, Sigma Aldrich, Cat# L1030 and L2778), diluted at a ratio of 1:5 in D-PBS. An image cytometer (BioTek Instruments Inc.) was used for imaging the networks.

### Vascular permeability and drug testing

To visualize the vascular permeability differences in the networks, 70kDa FITC dextran (Sigma Aldrich, Cat# 46945) were perfused through the networks after treatments and imaged using an image cytometer (BioTek Instruments Inc.). To quantify the permeability changes in the vascular networks that were exposed to various external stimulations, vessels were treated with Triton− X-100 (0.1% v/v), TNF-α (0.5 or 1 µg/mL, Sigma Aldrich, Cat# SRP3177) or lipopolysaccharides (LPS) from Escherichia coli (200 µg/mL, Sigma Aldrich, Cat# L4391) for 24 hours before the permeability test. 70kDa FITC dextran were perfused through the networks after each treatment. The fluorescent intensities in the middle wells were measured 0 minute and 15 minutes after adding the dextran solution. The change in fluorescent intensity was quantified and correlated to a standard curve to calculate the amounts of dextran diffusing across the vascular barrier.

### Kidney tubule permeability

To assess the permeability of the kidney proximal tubules and vessels, we perfused the networks with 70kDa FITC and TRITC-labelled dextran (Sigma Aldrich, Cat# T1162). We added 90 µL of FITC and TRITC-labelled dextran (500 µg/mL) in D-PBS to the inlet well of the vessel and proximal tubules respectively and 60 µL of D-PBS to the tissue well. To visualize perfusion of the dextran molecules through the tubular and vascular network, we used an image cytometer (BioTek Instruments Inc.). From the time-lapse perfusion images taken at 15-minute intervals, we calculated the diffusive permeability of the vascular and tubular network, P_d_, as previously reported^20,21^. I_i_ and I_f_ are the average intensities at final and initial timepoint and I_b_ is the average background intensities. Δt is the time interval and d is the average diameter of the tubule or vessel channels.

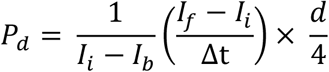

### Glucose reabsorption assay

To model glucose reabsorption carried out by the proximal tubules, we added ProxUp and ECGM2 media mixed at 1:1 ratio and containing an additional 5.5 mmol/L of D-Glucose (Sigma Aldrich, Cat# G5767) to all the wells on days 5, 9 and 13. Media perfusates from the tubular and vascular networks were collected separately after 24 hours and glucose levels were measured using a calibrated glucometer (Contour^®^ Next One meter).

### Cytokine assay

To stimulate renal tissue injury, culture media containing 50 ng/mL of TNF-α were added to the vascular network and incubated at 37 °C for 12 hours. Tissues incubated with culture media not supplemented with TNF-α were used as control. After 12 hours, media perfusates from treated and untreated tubular and vascular channels were collected separately. The collected perfusates were then centrifuged at 1000 rpm to remove cell debris and the supernatants were collected and stored at −80 °C until analysis. The collected supernatant was sent to Eve Technologies for quantifying pro-inflammatory cytokines using Human Cytokine Array Proinflammatory Focused 15-plex panel (Cat# HDF15). The tissues were also fixed and stained for ICAM-1 and ICAM-1 expression was quantified using ImageJ from the fluorescent images.

### Fabrication of AngioPlate™ lid for mechanical actuation

A SU8 master mold patterned with air distribution channels were made with standard photolithography technique (See **Supplementary Figure 6** for specific design dimensions). PDMS was mixed at a ratio of 1:10 and poured onto the SU8 master mold and left to cure at room temperature for three days. The PDMS sheet was then de-molded and trimmed to match the dimensions of a standard 384-well plate. An array of 2 mm holes was punched in the channel outlets using a biopsy punch (VWR, Cat# 21909-132). A hole was drilled on a plate lid (VWR, Cat# 10814-226) for the air inlet. 10-15 g of PDMS at a ratio of 1:10 ratio was then poured onto the lid. The drilled hole was temporarily blocked with a 1000 μL pipette tip (Fisherbrand, Cat# 02-707-507). To make O-rings, approximately 10-15 g of PDMS at 1:30 ratio was poured onto a one-well plate and left to cure for two days at room temperature. The 40 O-rings were made from the PDMS sheet with a 10 mm puncher (VWR, Cat# CA-95039-098) for the outer perimeter and a 2 mm biopsy puncher for the inner perimeter. The PDMS sheet with the air distribution channels and the plate lid with PDMS cover were both plasma-treated using an Electro-Technic Products Corona Plasma treater (Model BD-20AC). They were then bonded to each other. Next, the O-rings were aligned to the channel outlets and bonded to the assembled plate lid with plasma treatment. A silicon tubing with an outer diameter of 4 mm was then glued to the air inlet hole on the plate lid. The tubing is then connected to a syringe filter, another tubing extension, and then lastly, an external ventilator (Harvard Apparatus, Model 683 Volume Controlled Small Animal Ventilator, Cat# 55-0000). Before mechanical actuation, the plate lid was sterilized using 70% ethanol. The plate lid is secured onto the well plate with three rubber bands to ensure an air-tight seal. The tissues were actuated at 2.5 cc tidal volume and 30 breathes/min. The actuation process was visualized using a Nikon brightfield microscope and analyzed from the recorded video using the PIV plugin in ImageJ.

### Albumin assay

To evaluate the vascularized hepatic tissue, albumin production was analyzed as the primary marker. Culture media from inlet and outlet wells were collected on days 12, 16, and 20 and were analyzed with the Bethyl’s human albumin ELISA quantification Set (Bethyl Laboratories, Inc., Cat# E80-129) following the manufacturer’s instruction.

### Photoacoustic imaging

Ultrasound imaging was performed using the Vevo LAZR-X multi-modal *in vivo* imaging platform equipped with a MX 400 transducer (50 µm of axial resolution). A transparent bottomless 24-well plate was used to make the device for this application to minimize background noise. Before imaging, the vascular tissue was perfused with IR-783 dye (50 µM, Sigma-Aldrich, Cat# 543292-250MG), and the well with the tissue was filled with ultrasound gel. By controlling the movement of the transducer, the signal from the contrast agent was captured and recorded.

### Immunostaining and histology

To prepare for immunostaining, culture media was first removed from all wells and the tissues were washed with D-PBS three times. The tissues were fixed in 4% paraformaldehyde solution (Electron Microscopy Sciences, Cat# EMS 15710-S) at 4 °C overnight. Fixed tissues were washed with D-PBS, then blocked and permeated overnight at 4 °C under perfusion with 10% FBS containing 0.1% Triton™ X-100. Then the tissues were incubated in primary antibody solution for 48 hours at 4 °C followed by D-PBS wash at 4 °C for 48 hours to remove residual primary antibodies from the gel. Secondary or conjugated antibodies were added along with DAPI and incubated for 48 hours at 4 °C. The tissues were then washed with D-PBS for 48 hours at 4 °C prior to imaging with image cytometer. Dilutions used for all antibodies are shown in the table below. For 3D confocal images, stained tissues were imaged using Leica SP5 confocal microscopy. For histology sectioning, tissues were fixed in 10% formalin solution for 48 hours before being processed at the Histology Facility of McMaster University. The transmission electron microscope images showing the subcellular features of the endothelium were imaged with the Transmission Electron Microscope at the Electron Microscopy Facility of the Canadian Centre for Electron Microscopy at McMaster University. Antibody used and Catalogue #

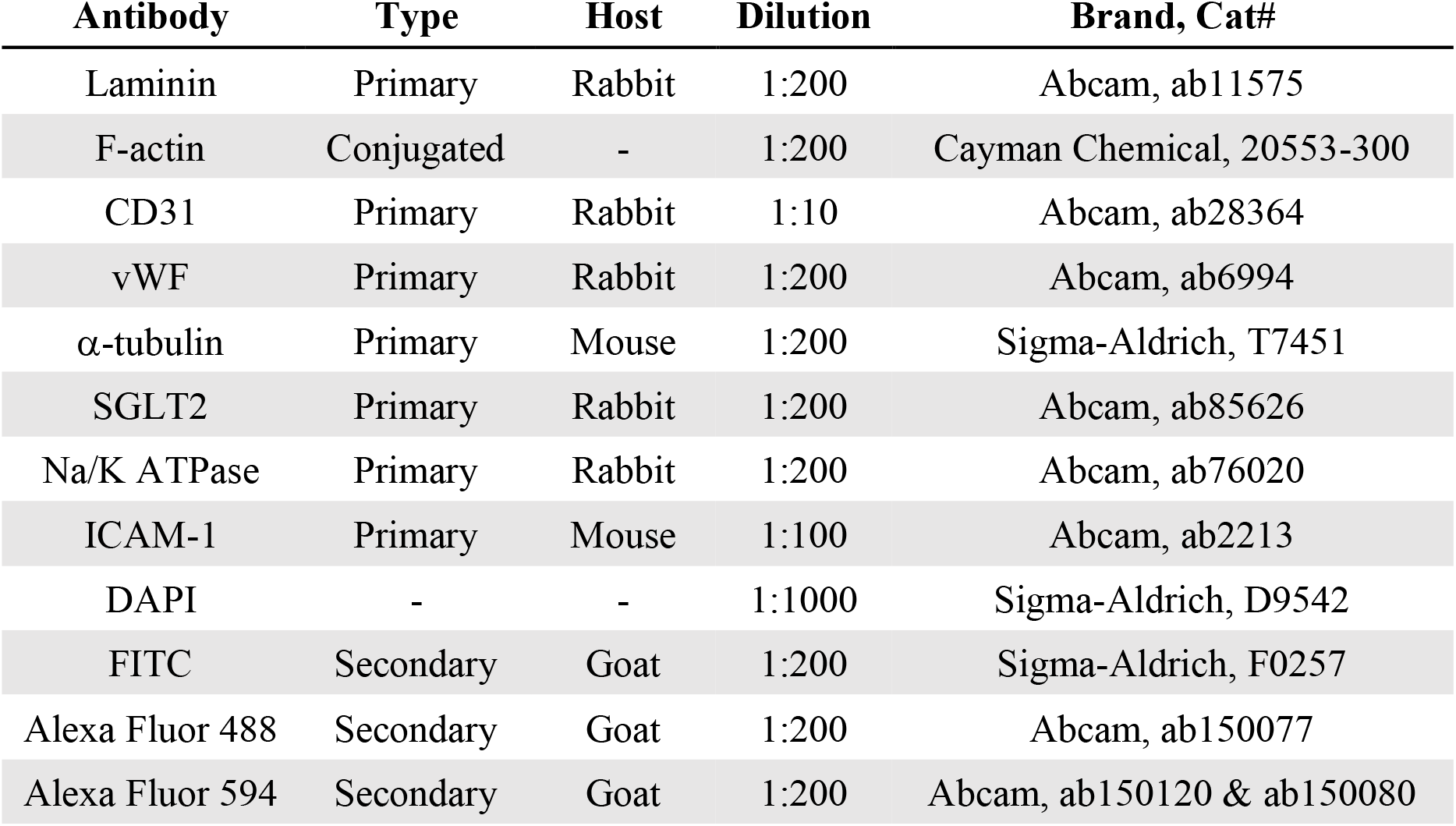

| **Antibody** | **Type** | **Host** | **Dilution** | **Brand, Cat#** |
| --- | --- | --- | --- | --- |
| Laminin | Primary | Rabbit | 1:200 | Abcam, ab11575 |
| F-actin | Conjugated | - | 1:200 | Cayman Chemical, 20553-300 |
| CD31 | Primary | Rabbit | 1:10 | Abcam, ab28364 |
| vWF | Primary | Rabbit | 1:200 | Abcam, ab6994 |
| α-tubulin | Primary | Mouse | 1:200 | Sigma-Aldrich, T7451 |
| SGLT2 | Primary | Rabbit | 1:200 | Abcam, ab85626 |
| Na/K ATPase | Primary | Rabbit | 1:200 | Abcam, ab76020 |
| ICAM-1 | Primary | Mouse | 1:100 | Abcam, ab2213 |
| DAPI | - | - | 1:1000 | Sigma-Aldrich, D9542 |
| FITC | Secondary | Goat | 1:200 | Sigma-Aldrich, F0257 |
| Alexa Fluor 488 | Secondary | Goat | 1:200 | Abcam, ab150077 |
| Alexa Fluor 594 | Secondary | Goat | 1:200 | Abcam, ab150120 & ab150080 |

### Statistical analysis

SigmaPlot and Prism were used for all statistical analyses. All data were tested for normality and equality of variance. To determine statistical significance, one-way ANOVA or one-way ANOVA on ranks with the Holm-Sidak method was applied. Data in all graphs were plotted as mean with standard deviation using Graphpad and at least three independent samples were used per condition for all quantitative analysis.

## Acknowledgments

We thank Dr. Samantha Slikboer for her help in performing ultrasound imaging and data processing. We thank Sonya Kouthouridis for her help in conducting histology of lung and kidney tissues. This work was made possible by the financial support of the National Sciences and Engineering Research Council of Canada (NSERC) CGS-D scholarship to S.R and D.S.Y.L. This work was also funded by the Canadian Institute of Health Research (CIHR) Project Grant (PJT-166052) to B.Z. We also would like to acknowledge BioRender.com for its support in creating figure illustrations.

## Author contribution

S.R., D.S.Y.L., and F.Z., performed the experiments, analyzed the results, and prepared the manuscript. A.S designed and fabricated the lung mechanical actuation plate lid. A.B., S.C. performed the Galectin-1 assay. A.L contributed to device fabrication. J.H., edited the manuscript. S.O. performed the albumin assay and edited the manuscript. A.K., edited the manuscript. B.Z. envisioned the concept, performed the experiments, supervised the work and edited the manuscript.

## Table of Contents

Inspired by 4D bioprinting, we develop a 4D subtractive manufacturing technique where a flexible sacrificial material can be patterned on a 2D surface, trigger to change shape when exposed to aqueous hydrogel, and subsequently degrade to produce complex perfusable networks in a natural extracellular matrix that can be populated with cells. The technique is applied to fabricate organ-specific vascular networks, vascularized kidney proximal tubule, lung alveoli terminal in a 384-well plate format, which is further scaled to a 24-well plate format to make a large vascular network, vascularized liver tissues, and for ultrasound imaging.

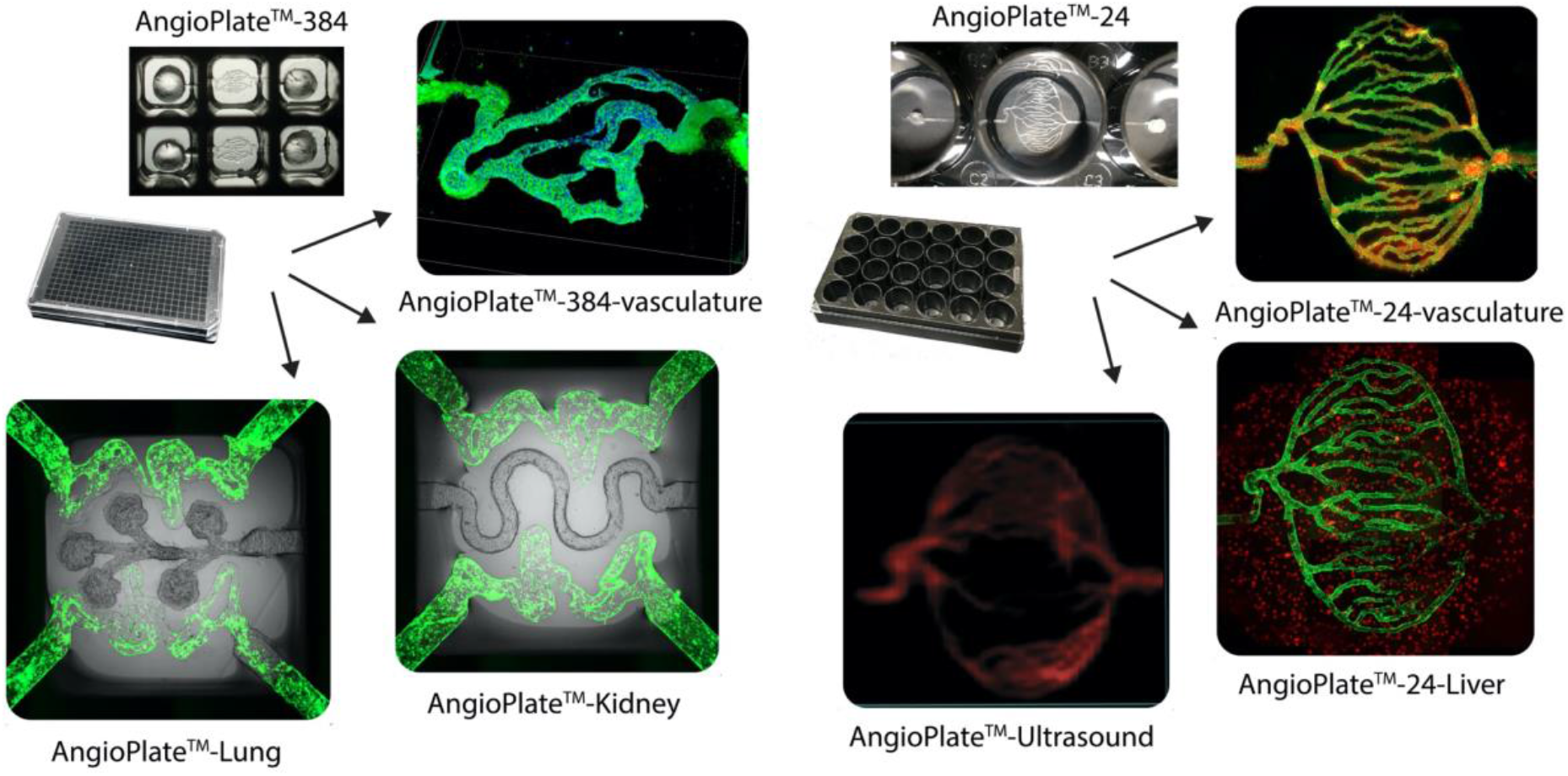

## Supplementary Materials

**Supplementary Figure 1.**
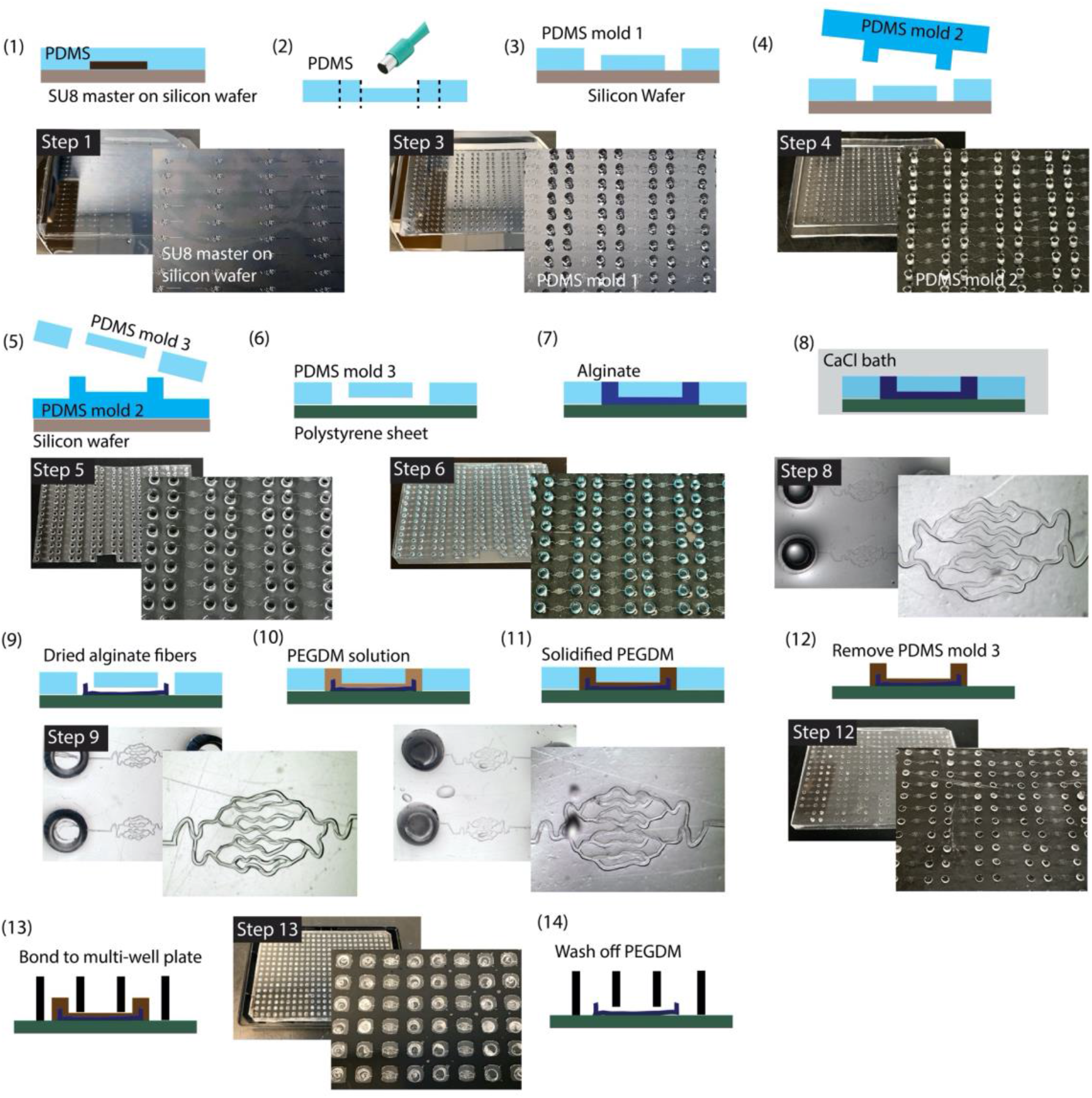
Step-by-step fabrication of AngioPlate™-384. First, using standard photolithography, we fabricated a PDMS mold with various patterns connected to an inlet and outlet well. The mold was then capped onto a polystyrene sheet to form an array of micro-channel networks. The networks were loaded with 3 wt.% alginate solution. Next, the entire mold was immersed in a calcium bath (1 mM), where calcium ions gradually diffused from the inlet and outlet wells into the alginate solution within the network, cross-linking the alginate overnight. With this approach, we were able to pattern 128 independent alginate fiber networks in the format of a 384-well plate. PEGDM solution was injected into the channels in the same way to encapsulate the alginate fiber to facilitate alginate release and to create the inlet/outlet channels. Finally, the polystyrene sheet patterned with alginate and PEGDM was assembled onto the base of a bottomless 384-well plate, encasing and sealing the alginate networks with a high viscosity PDMS glue.

**Supplementary Figure 2.**
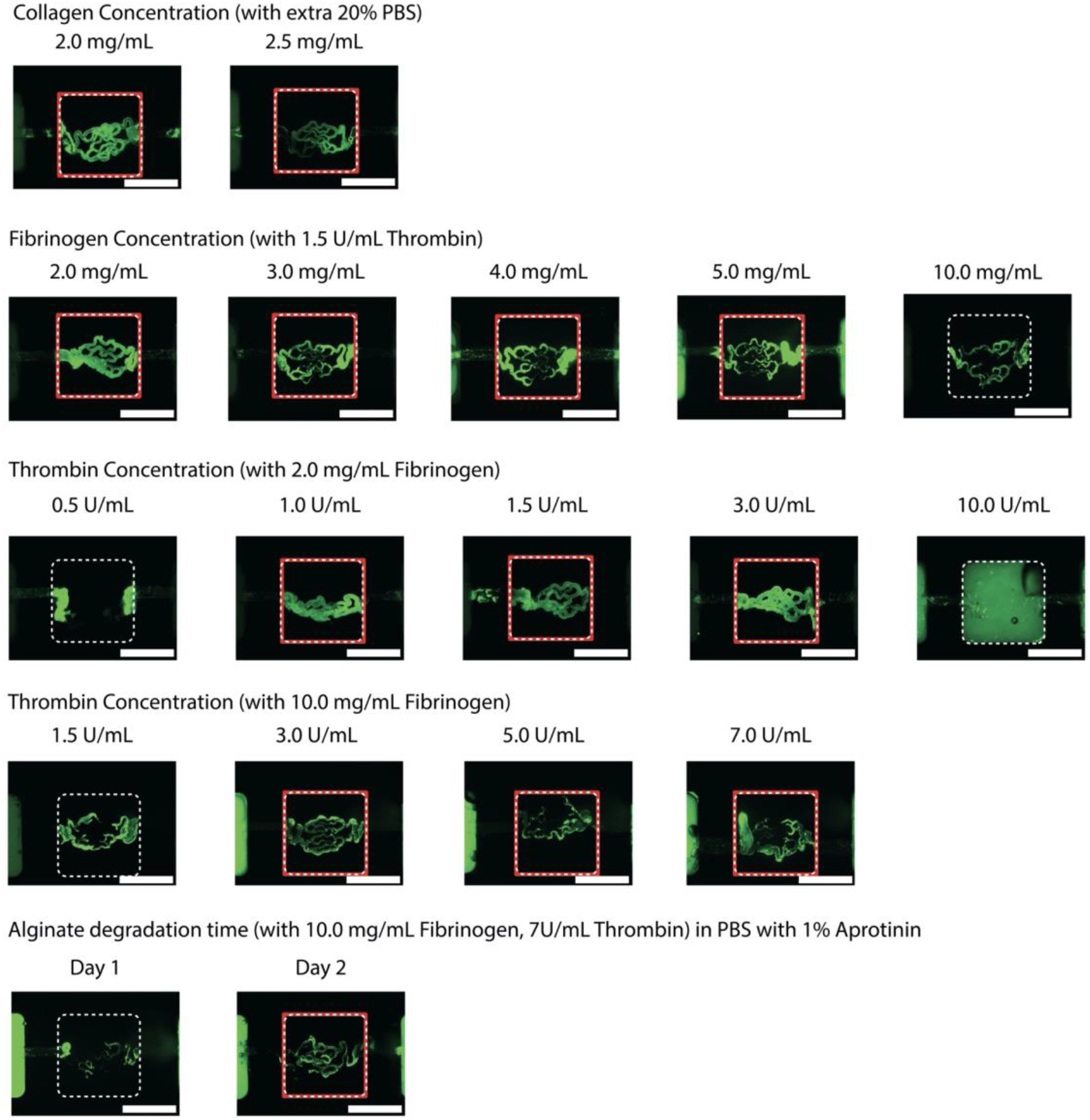
Optimization of hydrogel matrix cross-linking condition for network formation. Fluorescent images of networks perfused with 1 µm fluorescent particles (green) under various gelling conditions in both collagen-based gel and fibrin-based gel. Red boxes label the good conditions that resulted in the formation of complete perfusable networks. Scale bar, 2mm.

**Supplementary Figure 3.**
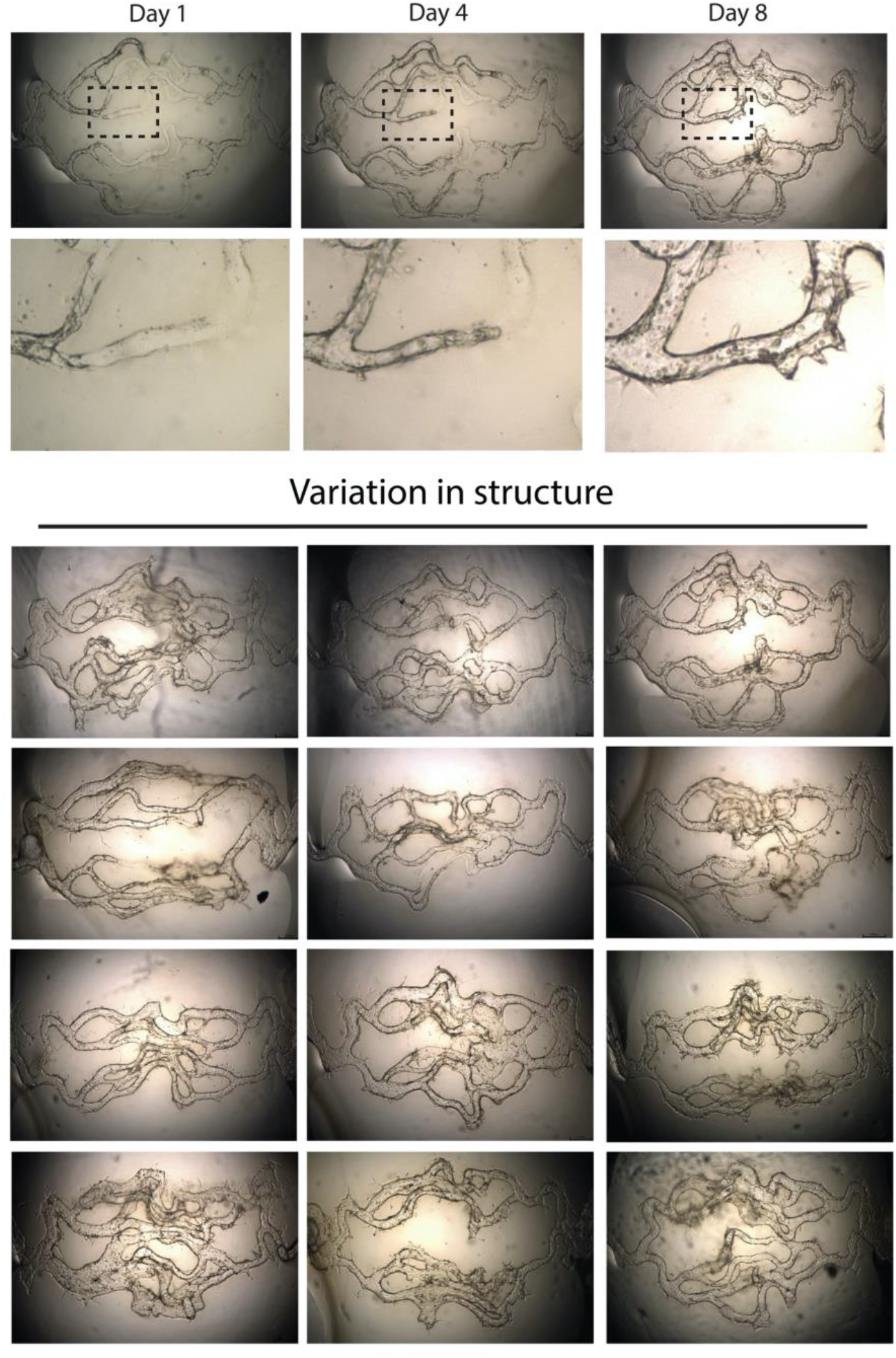
Structural variation in AngioPlate™-384-vasculature. Brightfield time-lapse images of networks seeded with human endothelial cells and variation in the network structures from 12 different wells.

**Supplementary Figure 4.**
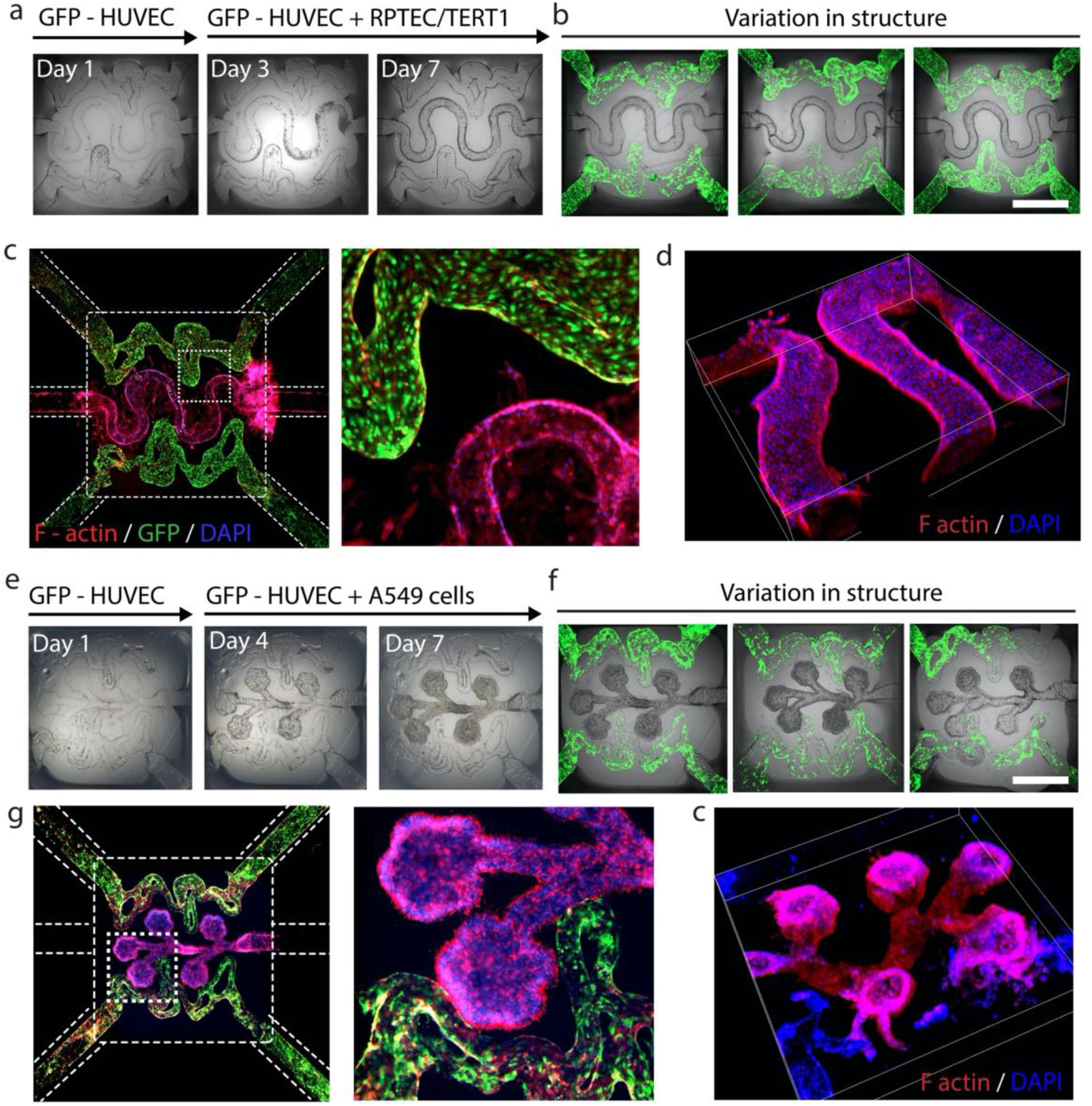
AngioPlate™-Lung and AngioPlate™-Kidney. **a**, Time-lapse brightfield images of a vascularized proximal tubule complex. Endothelial cells were seeded on day 0 and tubular cells were seeded on day 3. **b**, Variation in the structure of the vascular proximal tubule complex resulting from alginate folding. Scale bar 1 mm. **c-d**, Fluorescent images of vascular proximal tubule complex stained for F-actin (red) and DAPI (blue). **e**, Time-lapse brightfield images of a vascularized alveoli terminal. Endothelial cells were seeded on day 0 and Alveolar cells were seeded on day 3. **f**, Variation in the structure of vascularized alveoli terminal resulting from alginate folding. Scale bar 1 mm. **g-h**, Fluorescent images of a vascularized alveoli terminal stained for F-actin (red) and DAPI (blue).

**Supplementary Figure 5.**
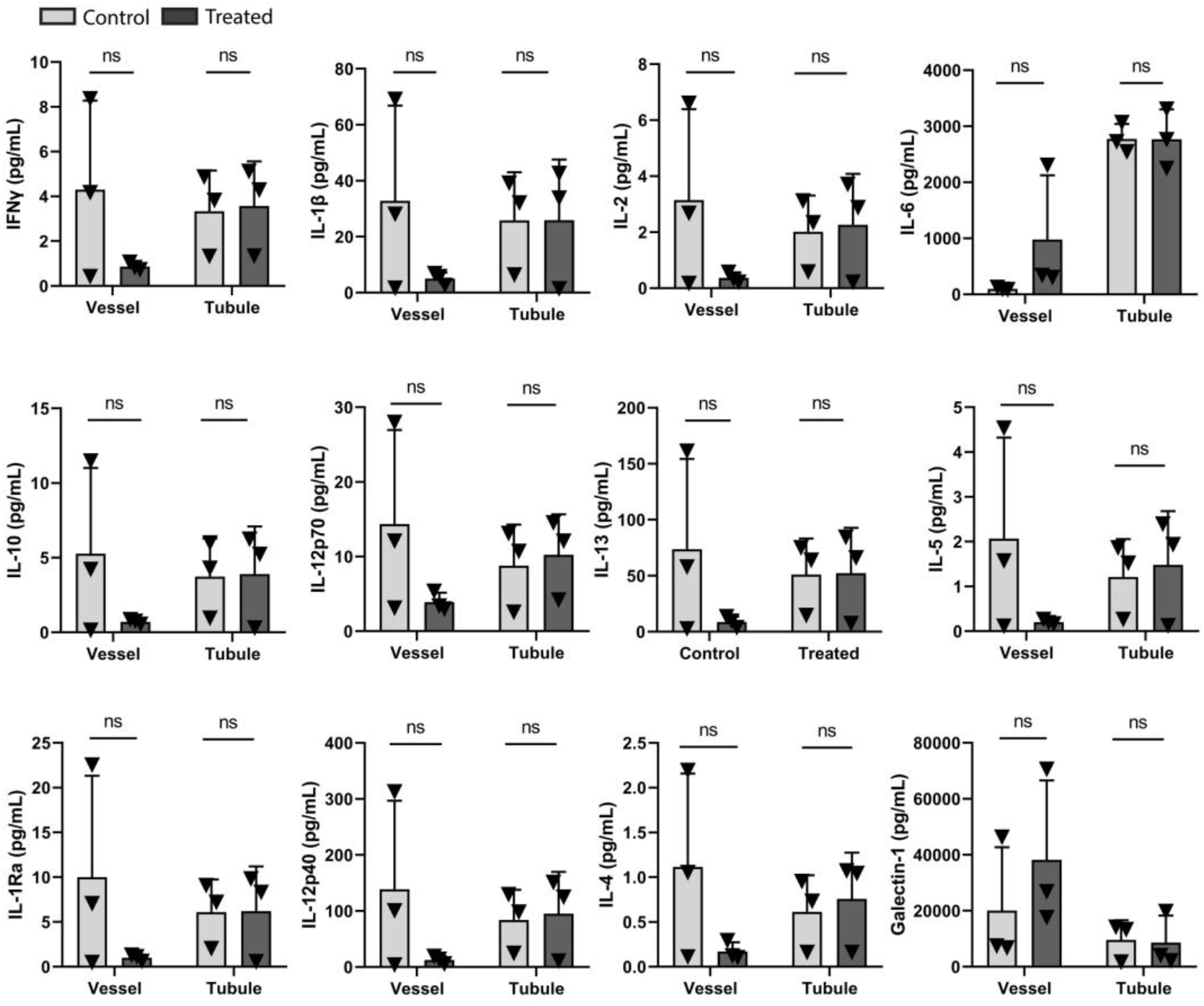
Cytokine assay of inflammatory AngioPlate™-Kidney. Cytokine analysis of media perfusates from tissues treated with or without TNF-α, n=3. Statistical significance was determined using one-way ANOVA and one-way ANOVA on ranks with the Holm-Sidak method. ns indicates not significantly different. *P≤0.05 **P≤0.01 ***P ≤0.001

**Supplementary Figure 6.**
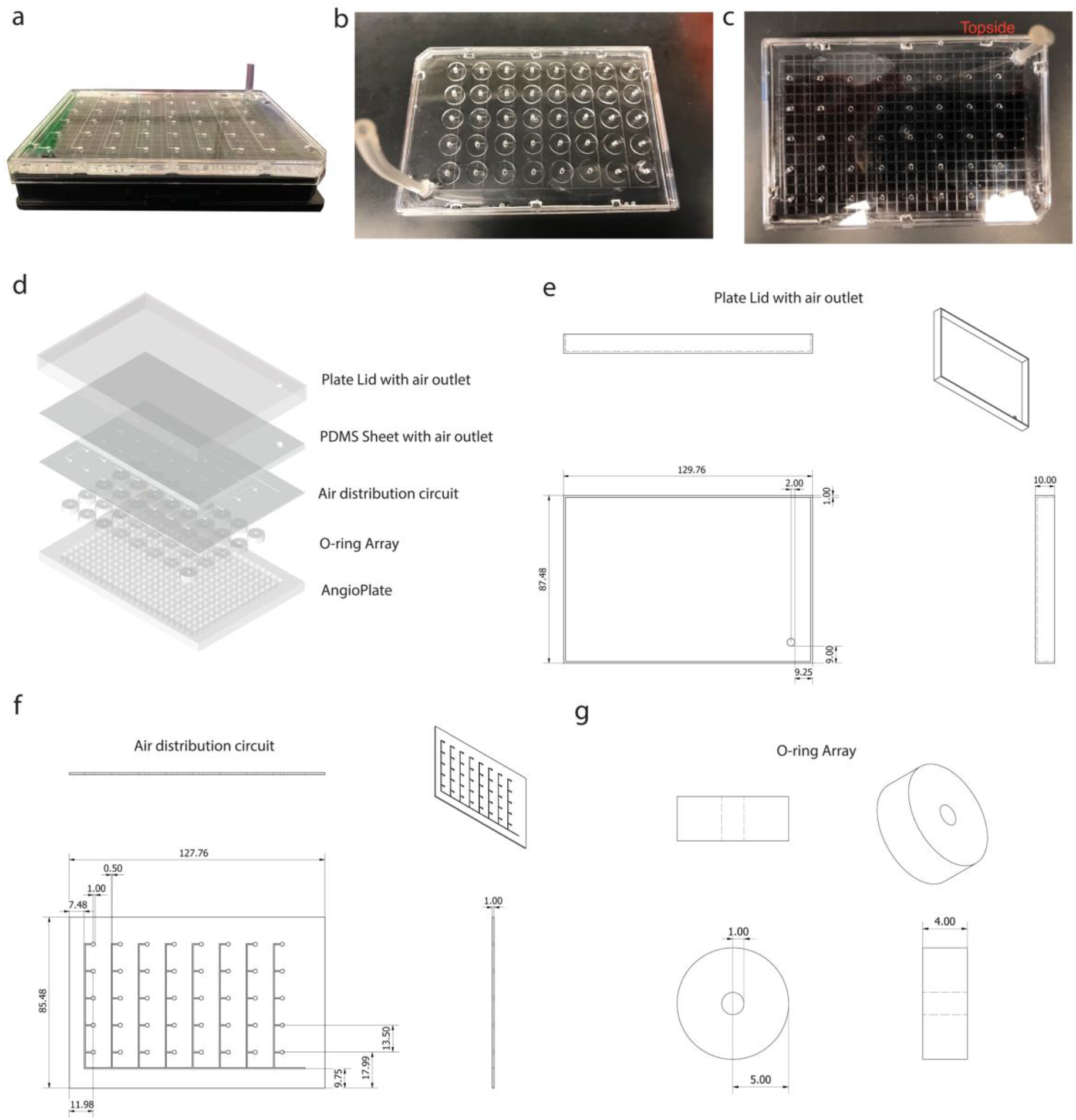
Mechanical actuation lid design, dimension and assembly. **a**, Image of the plate lid on an AngioPlate™. **b**, Top-down view of the actuation lid. **c** Top-down view of the actuation lid on the AngioPlate™ without the O-ring array to show alignment with the wells in an AngioPlate™. **d**, Assembly of different parts in the actuation lid. **e-g**, Dimensions and design of parts of the plate actuation lid, including the air distribution circuit and the o-rings. All labels are shown in millimeters.

**Supplementary Video 1**. Shape changing alginate fibers on AngioPlate

**Supplementary Video 2**. Mechanical actuation of terminal alveoli on AngioPlate

## References

1 Lee, A. et al. 3D bioprinting of collagen to rebuild components of the human heart. Science 365, 482–487 (2019).

2 Grigoryan, B. et al. Multivascular networks and functional intravascular topologies within biocompatible hydrogels. Science 364, 458–464 (2019).

3 Kinstlinger, I. S. et al. Perfusion and endothelialization of engineered tissues with patterned vascular networks. Nature Protocols 16, 3089–3113, doi:10.1038/s41596-021-00533-1 (2021).

4 Novak, R. et al. Robotic fluidic coupling and interrogation of multiple vascularized organ chips. Nature Biomedical Engineering 4, 407–420, doi:10.1038/s41551-019-0497-x (2020).

5 Trietsch, S. J. et al. Membrane-free culture and real-time barrier integrity assessment of perfused intestinal epithelium tubes. Nature Communications 8, 262 (2017).

6 Lin, N. Y. C. et al. Renal reabsorption in 3D vascularized proximal tubule models. Proceedings of the National Academy of Sciences 116, 5399–5404, doi:10.1073/pnas.1815208116 (2019).

7 Brassard, J. A., Nikolaev, M., Hübscher, T., Hofer, M. & Lutolf, M. P. Recapitulating macro-scale tissue self-organization through organoid bioprinting. Nature Materials, doi:10.1038/s41563-020-00803-5 (2020).

8 Nikolaev, M. et al. Homeostatic mini-intestines through scaffold-guided organoid morphogenesis. Nature, doi:10.1038/s41586-020-2724-8 (2020).

9 Arakawa, C. et al. Biophysical and biomolecular interactions of malaria-infected erythrocytes in engineered human capillaries. Science Advances 6, eaay7243 (2020).

10 Rayner, S. G. et al. Multiphoton-Guided Creation of Complex Organ-Specific Microvasculature. Advanced Healthcare Materials 10, 2100031, doi:https://doi.org/10.1002/adhm.202100031 (2021).

11 Gladman, A. S., Matsumoto, E. A., Nuzzo, R. G., Mahadevan, L. & Lewis, J. A. Biomimetic 4D printing. Nature materials (2016).

12 Leng, L., McAllister, A., Zhang, B., Radisic, M. & Günther, A. Mosaic Hydrogels: One-Step Formation of Multiscale Soft Materials. Adv. Mater. 24, 3650–3658 (2012).

13 Tønnesen, H. H. & Karlsen, J. Alginate in drug delivery systems. Drug development and industrial pharmacy 28, 621–630 (2002).

14 Kouthouridis, S. et al. Oxygenation as a driving factor in epithelial differentiation at the air–liquid interface. Integrative Biology 13, 61–72, doi:10.1093/intbio/zyab002 (2021).

15 Samal, P., van Blitterswijk, C., Truckenmüller, R. & Giselbrecht, S. Grow with the Flow: When Morphogenesis Meets Microfluidics. Adv. Mater. 31, 1805764 (2019).

16 Campbell, S. B. et al. Beyond Polydimethylsiloxane: Alternative Materials for Fabrication of Organ-on-a-Chip Devices and Microphysiological Systems. ACS Biomaterials Science & Engineering (2020).

17 Azizgolshani, H. et al. High-throughput organ-on-chip platform with integrated programmable fluid flow and real-time sensing for complex tissue models in drug development workflows. Lab on a Chip 21, 1454–1474, doi:10.1039/D1LC00067E (2021).

18 Parrish, J., Lim, K., Zhang, B., Radisic, M. & Woodfield, T. B. F. New Frontiers for Biofabrication and Bioreactor Design in Microphysiological System Development. Trends in Biotechnology, doi:https://doi.org/10.1016/j.tibtech.2019.04.009 (2019).

19 Takebe, T., Zhang, B. & Radisic, M. Synergistic Engineering: Organoids Meet Organs-on-a-Chip. Cell Stem Cell 21, 297–300 (2017).

20 Jeon, J. S. et al. Generation of 3D functional microvascular networks with human mesenchymal stem cells in microfluidic systems. Integr. Biol. 6, 555–563, doi:10.1039/c3ib40267c (2014).

21 Sobrino, A. et al. 3D microtumors in vitro supported by perfused vascular networks. Scientific Reports 6, 31589, doi:10.1038/srep31589 (2016).

